# Non-standard viral genome-derived RNA activates TLR3 and type I IFN signaling to induce cDC1-dependent CD8+ T-cell responses during vaccination in mice

**DOI:** 10.1101/2022.06.09.495494

**Authors:** Devin G. Fisher, David J. Holthausen, Carolina B. López

**Author notes:** Corresponding author: Dr. Carolina B. López, Department of Molecular Microbiology, Washington University School of Medicine, St. Louis, MO.

## Abstract

There is a critical need to develop vaccine adjuvants that induce robust immune responses able to protect against intracellular pathogens, including viruses. Previously, we described the virus-derived adjuvant, defective viral genome-derived oligonucleotide (DDO), as a strong inducer of type 1 immune responses, including protective Th1 CD4+ T-cells and effector CD8+ T-cells in mice. Here we unravel the early innate response required for this type 1 immunity induction. Upon DDO subcutaneous injection, type 1 conventional dendritic cells (cDC1s) accumulate rapidly in the draining lymph node in a type I interferon (IFN)-dependent manner. cDC1 accumulation in the lymph node is required for antigen-specific CD8+ T-cell responses. Notably, in contrast to poly I:C, DDO administration resulted in type I IFN expression at the injection site, but not in the draining lymph node. Additionally, DDO induced an inflammatory cytokine profile distinct from that induced by poly I:C. Therefore, DDO represent a powerful new adjuvant to be used during vaccination against intracellular pathogens.

**IMPORTANCE:** There is a paucity of vaccine adjuvants able to trigger effective and safe protective responses to many intracellular pathogens. Defining the minimal requirements to achieve type 1 immunity, which includes antigen specific CD8+ T cells capable of eliminating infected cells, is essential for the development of adjuvants that lead to optimal protective immune responses during vaccination against intracellular pathogens. We use a virus-derived immunostimulatory molecule, defective viral genome-derived oligonucleotide (DDO), to provide insights into how type 1 immune responses are triggered during vaccination using an inactivated influenza vaccine model. Understanding the mechanism of action of vaccine adjuvants not only aids in the advancement of vaccine development, but also in understanding specific immune pathways required for efficient induction of adaptive immune responses to infections.

## 1. INTRODUCTION

Over the last two centuries, vaccines have saved countless lives[1]. However, recent epidemics and pandemics have highlighted the continued need for vaccines against newly emerging pathogens. The development of new vaccines is constrained by safety issues and by our ability to tailor vaccines to induce the right type of immune response capable of controlling and eliminating the target pathogen. Attenuated vaccines, while capable of inducing robust immune responses that mimic those generated during natural infections, present challenges due to their ability to revert to virulence or cause disease in a growing population of immunocompromised people[2]. Messenger RNA (mRNA) vaccines have been instrumental in efforts to quelle the COVID-19 pandemic, however the extent of lasting immunity remains has yet to be determined [3]. Killed or subunit vaccines offer a safer alternative to attenuated vaccines, however, they require an adjuvant to induce protective immune responses. Adjuvants not only provide a necessary stimulus for the immune response to vaccinated antigens, but also shape the induced immune response[4].

Type 1 immunity, including a robust CD8+ T-cell response, is critical for clearance of many intracellular pathogens. However, currently we lack adjuvants capable of inducing robust CD8+ T-cell responses thereby limiting our ability to develop effective vaccines against many pathogens. Type 2 immunity is commonly induced by existing adjuvants and consists of antibodies and Th2 CD4+ T-cells. While the optimal response for extracellular pathogens, this type of response can be detrimental to the host during many infections, especially from respiratory viruses. Early preclinical studies for SARS-CoV vaccines included alum, a type 2 immunity-inducing adjuvant, and resulted in worsened morbidity after challenge[5]. Similarly, type 2 immune responses induced upon respiratory syncytial virus (RSV) vaccination or infection led to worsened disease or even death[6-8]. Comparable findings were observed with rhinovirus infections in infants[9, 10]. These observations highlight the importance of developing adjuvants that induce type 1 immune responses upon vaccination against these classes of pathogens. However, to develop effective type 1 immunity-inducing adjuvants, we first must understand what early steps are required during vaccination that prompt a CD8+ T-cell response.

We have shown that a virus-derived RNA adjuvant named defective viral genome-derived oligonucleotide (DDO) induces a robust type 1 immune response in mice. This response synergizes with other licensed adjuvants and induces effector CD8+ T-cells leading to faster recovery from heterosubtypic influenza challenge[11]. DDOs are synthetic and replication incompetent RNAs derived from the 546nt Sendai virus (SeV) non-standard viral genome (NsVG) that is the primary immunostimulatory molecule during SeV infections[12, 13].

In this study, we asked how a 268 nt-long DDO was able to induce an antigen-specific CD8+ T-cell response. We investigated the type of dendritic cells (DCs) activated in response to DDO injection and if those DCs were critical for the CD8+ T-cell response. We next investigated the local cytokine response at the injection site and the draining lymph node to better understand the early response that leads to type 1 immunity. Finally, we uncovered how DDO-268 was sensed by cells to induce an immune response to DDO-adjuvanted vaccines. Together, this study helps build our understanding of how the early innate immune response to a type 1 immunity-inducing adjuvant leads to robust adaptive immunity.

## 2. METHODS

### 2.1 Ethics statement

All described studies carefully adhered to the recommendations in the Guide for the Care and Use of Laboratory Animal of the National Institute of Health. Institutional Animal Care and Use Committee, University of Pennsylvania Animal Welfare Assurance Number A3079-01 approved protocol 804691 and Washington University in St. Louis approved protocol 20-0120.

### 2.2 Mice

C57BL/6 mice were obtained from Jackson Laboratory. *Ifnar1*^-/-^ mice were a kind donation of Dr. Thomas Moran (Icahn School of Medicine at Mount Sinai)[14] and were used with sex and age matched C57BL/6 mice (Jackson Laboratory bred in house). *Mavs*^-/-^ mice (B6;129-Mavs^tm1Zjc^/J) and WT control recommended by Jackson labs (B6129SF2/J) were purchased from Jackson Laboratory. *Tlr3*^-/-^ mice (B6;129S1-Tlr3^tm1Flv^/J) and WT control recommended by Jackson laboratory (B6129SF2/J mice) were purchased from Jackson Laboratory. *Tlr7*^-^ (B6.129S1-Tlr7t^m1Flv^/J) and WT control recommended by Jackson laboratory (C57BL/6NJ) were purchased from Jackson Laboratory. *Batf3*^-/-^ mice were a kind gift from Dr. Christopher Hunter (School of Veterinary Medicine at the University of Pennsylvania). All mice used were 6 to 8 weeks old at the start of the experiment. All experiments were performed with male and female mice.

### 2.3 Vaccine formulation

Disrupted IAV (disIAV) vaccine: Influenza A/Puerto Rico/8/1934 H1N1 (IAV PR/8) was harvested 40h post-inoculation from the allantoic fluid of 10 day old embryonated eggs and purified through a 35% sucrose cushion. Virus was inactivated with UV light (254nm at 6-inch distance) for 40 minutes as previously described[11]. Inactivated virus was disrupted by freezing on dry ice and ethanol, thawing at 37°C, and vortexing for 1 minute. The freeze-thawing cycle was repeated three times. Inactivation was confirmed by the inability of the virus to replicate in MDCK cells (Madin-Darby canine kidney cells, gift from Dr. Scott Hensley, University of Pennsylvania) in the presence of 2mg/ml trypsin. DDO is a single stranded 268nt *in vitro*-transcribed RNA that contains the immunostimulatory motif of a DVG and was previously characterized[15]. DDO was produced, stored, characterized, and quality controlled as described[11, 12].

### 2.4 Mouse immunization and injection

For immunization, mice were anesthetized with isoflurane and injected subcutaneously (s.c.) into the rear footpad with 10µg disIAV vaccine diluted in PBS adjuvanted with 5µg DDO, 5µg Low Molecular Weight polyinosine-polycytidylic acid (poly I:C, InvivoGen), or Alum (Alhydrogel 2%, InvivoGen) at 50% v/v at final volume of 30µl per dose. Mice were primed and boosted 14 days later with the same vaccine formulation. For adjuvant only experiments, mice were anesthetized with isoflurane and injected s.c. into the rear footpad with PBS, 5µg DDO, or 5µg poly I:C diluted with PBS to a final volume of 30µl.

### 2.5 RT-qPCR from footpads and lymph nodes

The flesh of injected foot pads and draining lymph nodes were harvested at 4, 12, or 24h post-injection and placed in TRIzol (Invitrogen). Two lymph nodes were pooled to increase RNA yield and quality. One microgram of RNA isolated by TRIzol was reversed transcribed using high-capacity RNA to cDNA reagents (Applied Biosystem). qPCR assays were performed using SYBR Green PCR Master Mix (Applied Biosystem) in a Viia7 Applied Biosystem Lightcycler. Primers used in the assay were:

*B actin* for-5’-AGGTGACAGCATTGCTTCTG-3’ and rev-5’ GCTGCCTCAACACCTCAAC-3’, *Ifnb1* for-5’- AGATGTCCTCAACTGCTCTC-3’ and rev-5’- AGATTCACTACCAGTCCCAG-3’, *Mx1* for-5’- CAACTGGAATCCTCCTGGAA-3’ and rev-5’- GGCTCTCCTCAGAGGTATCA’3’, *Ccl5* for-5’- GCAGCAAGTGCTCCAATCTT-3’ and rev-5’- CAGGGAAGCGTATACAGGGT-3’, *Il6* for-5’- ACAGAAGGAGTGGCTAAGGA-3’ and rev-5’- CGCACTAGGTTTGCCGAGTA-3’, *Il1b* for-5’- TTGACGGACCCCAAAAGAT-3’ and rev-5’- GATGTGCTGCTGCGAGATT-3’, *Ifna1* for-5’- AGCCTTGACACTCCTGGTACA-3’ and rev-5’- TGAGCCTTCTTGATCTGCTG-3’, *Ifna2* for-5’- CCTGTGCTGCGAGATCTTACT-3’ and rev-5’- GGTGGAGGTCATTGCAGAAT-3’, *Ifna4* for-5’- AGCCTGTGTGATGCAGGAA-3’ and rev-5’- GGCACAGAGGCTGTGTTTCT-3’, *Ifna5* for-5’- CCTCAGGAACAAGAGAGCCTTA-3’ and rev-5’- TCCTGTGGGAATCCAAAGTC-3’, *Ifna6* for-5’- GATGGTTTTGGTGGTGTTGA-3’ and rev-5’- CCAGGAGTGTCAAGGCTTTC-3’, *Ifna12* for-5’- ACAGCCCAGAGGACAAACAG-3’ and rev-5’- GTCTGAGGCAGGTCACATCC-3’,, *Ifna13* for-5’- TGCTGGCTGTGAGGAAATACT-3’ and rev-5’- GGAAGACAGGGTTCTCTGGA-3’.

### 2.6 Flow cytometry

Single-cell suspensions of spleen and popliteal lymph node were prepared and stained with fluorochrome-labeled antibodies as previously described[16]. Popliteal lymph nodes used for DC staining were digested using DNase (1µg/ml) and Liberase (5µg/ml) in Roswell Park Memorial Institute (RPMI) 1640 Medium (Life Technologies) for 20min at 37°C. Fixable Viability Dye eFluor506, monoclonal antibody specific for mouse IFNγ (clone XMG1.2), Ly6c (clone HK1.4), CD3 (clone 17A2), CD19 (clone eBio1D3), B220 (clone RA3-6B2), NK1.1 (clone PK136), CD11b (clone M1/70), and TNFα (clone MP6-XT22) were obtained from eBioscience. Monoclonal antibodies specific for mouse CD3 (clone 17A2), CD4 (clone GK1.5), CD11a (clone H11578), XCR1 (clone ZET), PDCA1 (clone 129cl), CD11c (clone N418), SIRPα (clone P84), CD64 (clone x54/7.1) and MHC-II (clone M5/114.15.2) were obtained from Biolegend. Monoclonal antibodies specific for mouse CD86 (clone GL1) and CD8 (clone 53-6.7) were obtained from BD BioSciences. IAV-specific tetramers: H-2D^b^ tetramers bearing NP366-374 (ASNENMETM) and I-A^b^ tetramers bearing NP311-325 (QVYSLIRPNENPAHK) were obtained from NIH Tetramer Core Facility at Emory University. Samples were acquired on a LSRFortessa (BD Bioscience) cytometer and analyzed using the FlowJo Software (TreeStar).

### 2.9 Statistical analysis

Statistical analysis was performed using GraphPad Prism 8 and 9 for Mac (GraphPad Software). RT-qPCR and flow cytometry data were analyzed by two-way ANOVA followed by Tukey’s multiple comparison test.

## 3. RESULTS

### 3.1. Subcutaneous injection of a DDO-adjuvanted vaccine induces CD8+ T-cell responses

We previously showed that upon intramuscular immunization, DDO-268 (DDO from now on) induces a robust type 1 immune response in mice that included antigen-specific IgG2c antibodies, TNFα- and IFNγ-producing CD4+ T-cells, and CD8+ T-cells[11]. To confirm that DDO is effective in generating antigen-specific CD8+ T-cells when used during subcutaneous (s.c.) immunization, and to compare its activity with the gold standard RNA adjuvant dsRNA mimic polyinosine-polycytidylic acid (poly I:C), we used a model of disrupted influenza A virus (disIAV) vaccination. Mice were immunized twice, 14 days apart, with disIAV alone or adjuvanted with DDO, poly I:C, or the type 2 immunity-inducing adjuvant alum. To assess the T-cell response, spleens were harvested 7 days post-boost and antigen-specific T-cells were examined using IAV-specific tetramer staining. DDO induced greater antigen-specific CD8+ T-cell responses by both number and percentage than all other groups, including those immunized in the presence of poly I:C. Mice immunized with alum-adjuvanted vaccines completely failed to induce CD8+ T-cell responses, as expected (Fig 1).

**Figure 1:**
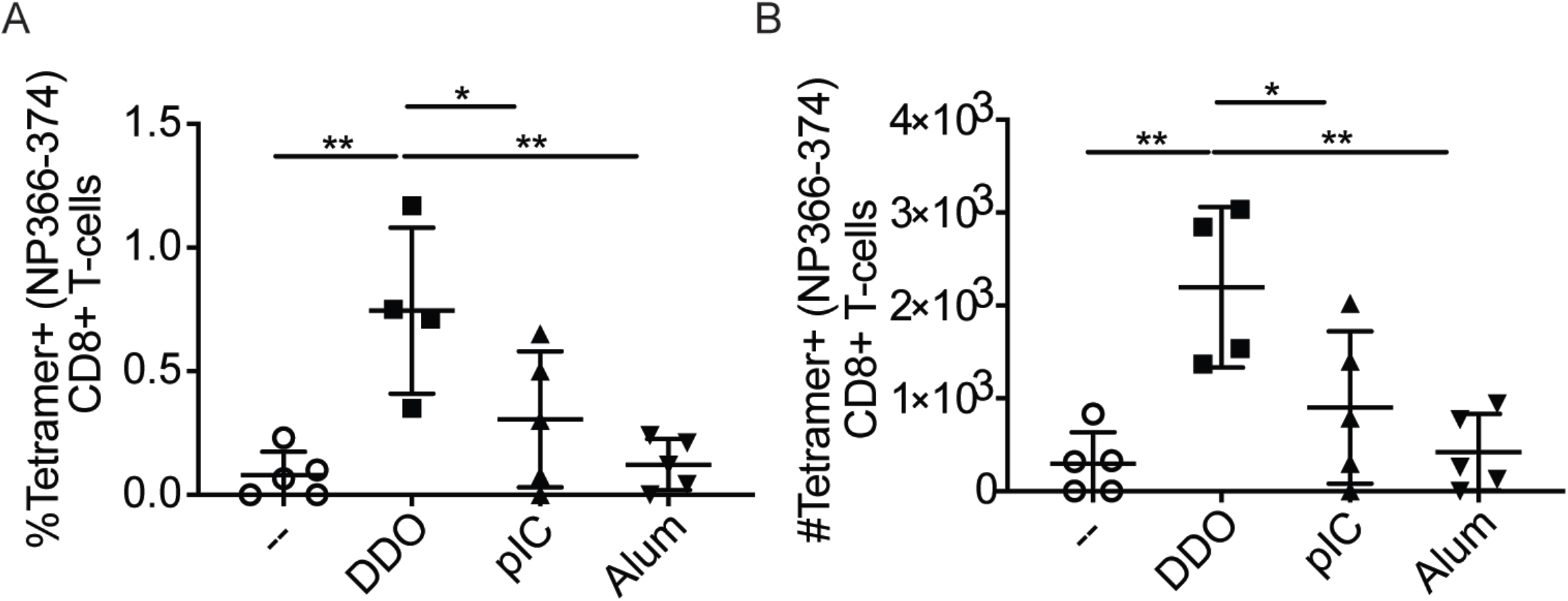
DDO induces a robust CD8+ T-cell response upon IAV immunization. WT mice were immunized twice, 14d apart, with 10µg inactivated and disrupted IAV PR/8 (disIAV) alone (--) or adjuvanted with 5µg DDO, 5µg pIC, or 50% volume Alum. (n=5 mice/group). **A-B** Spleens were harvested 7d post-boost and processed into a single cell suspension and examined by flow cytometry. Mean±SEM of each group is shown. Tetramer+ CD8+ T-cells were defined as live, CD3^+^, CD8^+^CD4^-^, tetramer^+^. **A** Percent of CD8+ T-cells that are tetramer+. (n=4-5 mice/group) **B** total number of antigen specific CD8+ T-cells in the spleen. *=p<0.05, **=p<0.01. Data represent one representative experiment out of 2 independent repeats.

### 3.2 DDO induces robust cDC1 accumulation in the draining lymph node

DCs are critical translators of the innate immune response into adaptive immunity[17]. DCs recruited to the draining lymph node at the time of T-cell priming direct and determine the adaptive immune response bias towards type 1 or type 2 immunity[18]. DCs are divided into three main subtypes, each with different functions. Type 1 conventional DCs (cDC1s) are specially equipped to take inert antigen, such as that found during killed or subunit vaccination, and cross-present it to activate CD8+ T-cells[18]. In contrast, cDC2s more readily activate CD4+ T-cells[19]. Finally, plasmacytoid DCs (pDCs) express high levels of virus sensors and are specially equipped to produce type I interferon (IFN) upon sensing viral nucleic acids but are not typically activators of naïve T-cells[20].

To examine the timing of DC recruitment to the draining lymph node and the changes that occurred in the DC composition of the draining lymph node upon s.c. injection, mice were injected with PBS, DDO, or poly I:C, and draining lymph nodes were harvested 12, 24, or 36h post-injection. At baseline, the conventional DC population composition in the lymph node is about 60% cDC2 (XCR1^-^, SIRPα^+^) and 40% cDC1 (XCR1^+^, SIRPα^-^) (Figure 2A, C). Both DDO and poly I:C injection increased the percentage and number of cDC1s in the lymph node which peaked at 12h post-injection (Figure 2A-B). cDC2s were reduced by percentage in the lymph node due to the early influx of cDC1s, with a maximal reduction in percentage at 12h post-injection (Figure 2C-D). The composition of the cDCs within the lymph node returned to baseline percentages by 36h post-injection, as shown by the ratio of cDC1 to cDC2 returning to the level of PBS (Figure 2E). In addition, we observed that the number and percentage of pDCs (B220^+^, PDCA1^+^) were increased in the lymph node reaching a maximum at 24hpi upon injection of both DDO and poly I:C (Figure 2F-G). Overall, s.c. injection of DDO induces robust and rapid accumulation of cDC1s and pDCs in the draining lymph node, similar to poly I:C.

**Figure 2:**
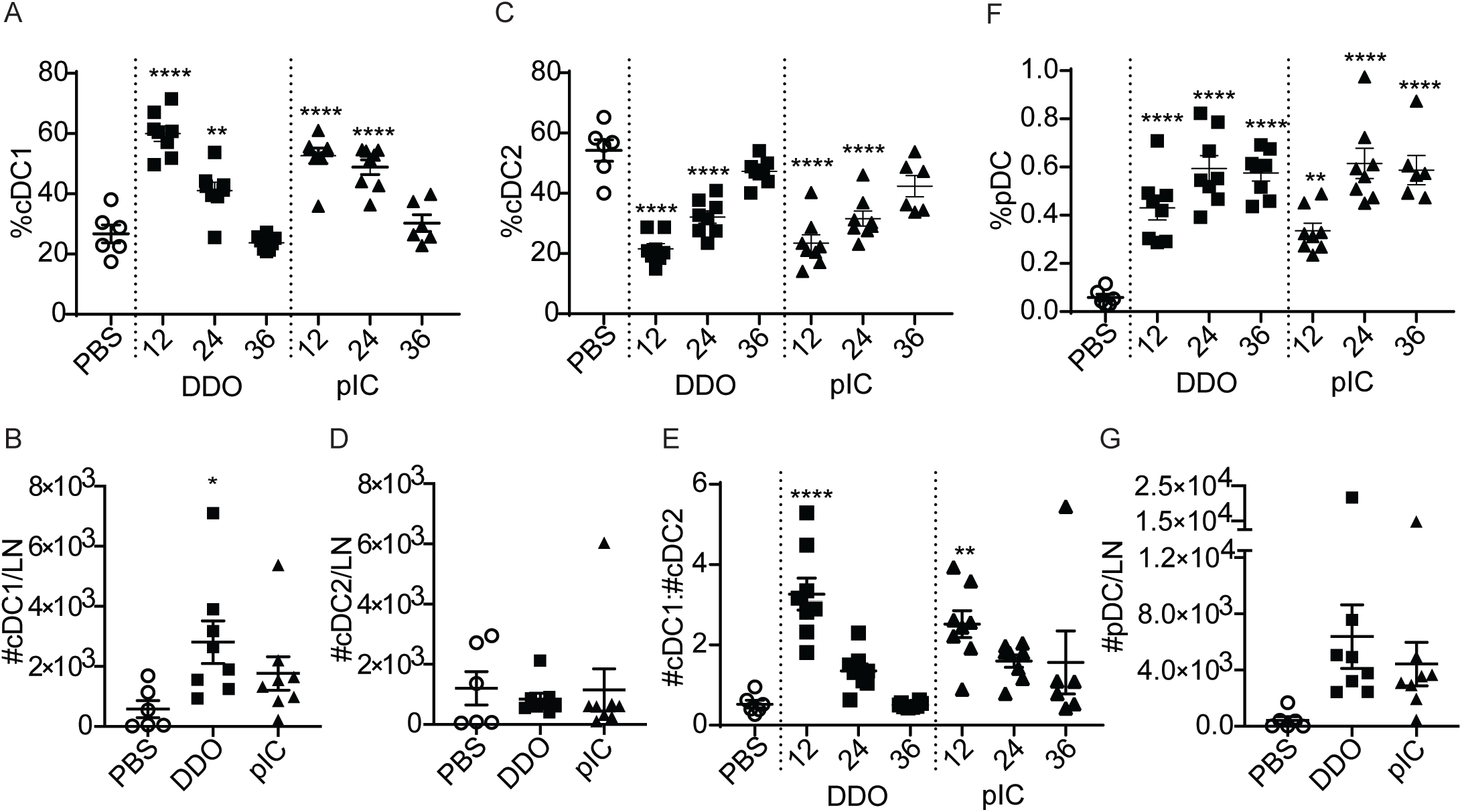
DDO induces accumulation of cDC1s in the draining lymph node. WT mice were injected s.c. with PBS, 5µg DDO, or 5µg poly I:C (pIC) in the rear footpad. Draining lymph node were collected 12, 24, or 36h post-injection for analysis. PBS injected mice were harvested 24h post-injection. (n= 6/group). Mean±SEM of each group is shown (**A-G**). **A-B** cDC1 were characterized as Live, CD3^-^NK1.1^-^B220^-^CD19^-^, MHCII^hi^, CD64^-^, Ly6c^-^, CD11c^hi^, XCR1^+^SIRPa^-^. **A** Percent of conventional DCs that are cDC1s. **B** Number of cDC1s in the draining lymph node at 12h post-injection. **C-D** cDC2 were characterized as Live, CD3^-^NK1.1^-^B220^-^CD19^-^, MHCII^hi^, CD64^-^,Ly6c^-^, CD11c^hi^, XCR1^-^SIRPa^+^. **C** Percent of conventional DCs that are cDC2s. **D** Number of cDC2s in the draining lymph node at 12h post-injection. **E** The ratio of cDC1 to cDC2 was created by dividing the number of cDC1 by number of cDC2 at the indicated time post-injection. **F-G** pDCs were characterized as live, PDCA1^+^B220^+^. **F** percent of cells in the lymph node that are pDCs. **G** Number of pDCs in the draining lymph node at 12h post-injection.*=p<0.05, **=p<0.01, ***=p<0.001, ****=p<0.0001 compared to PBS control. Data represent one representative experiment out of 3 independent repeats.

### 3.3 cDC1s are required for DDO-induced CD8+T-cell responses

cDC1s are excellent at cross-presenting antigen and are therefore important for the generation of CD8+ T-cell responses to subunit and killed vaccines[18]. Development of cDC1s, but not cDC2s or pDCs, depends on the transcription factor BATF3[21]. We first investigated the requirement for BATF3 in cDC1 accumulation in the draining lymph node after DDO injection. Mice lacking BATF3 (*Batf3*^-/-^) and WT mice were injected as described above and analysis of the composition of DCs in the lymph node 12h after-injection confirmed that BATF3 is required for cDC1 accumulation, but not for cDC2 or pDCs (Figure 3A-C). To assess the role of cDC1s in the induction of antigen-specific CD8+ T-cells responses to DDO-adjuvanted vaccines, mice lacking BATF3 and WT mice were vaccinated against IAV as described above. Spleens were harvested 7d post-boost for T-cell analysis. Antigen-specific CD8+ T-cells were only generated in response to DDO-adjuvanted vaccines in WT mice, but not in *Batf3^-/-^* mice lacking cDC1s (Figure 3D). Overall, these data show that DDO injection induced a cDC1 response that is required for antigen-specific CD8+ T-cell responses.

**Figure 3:**
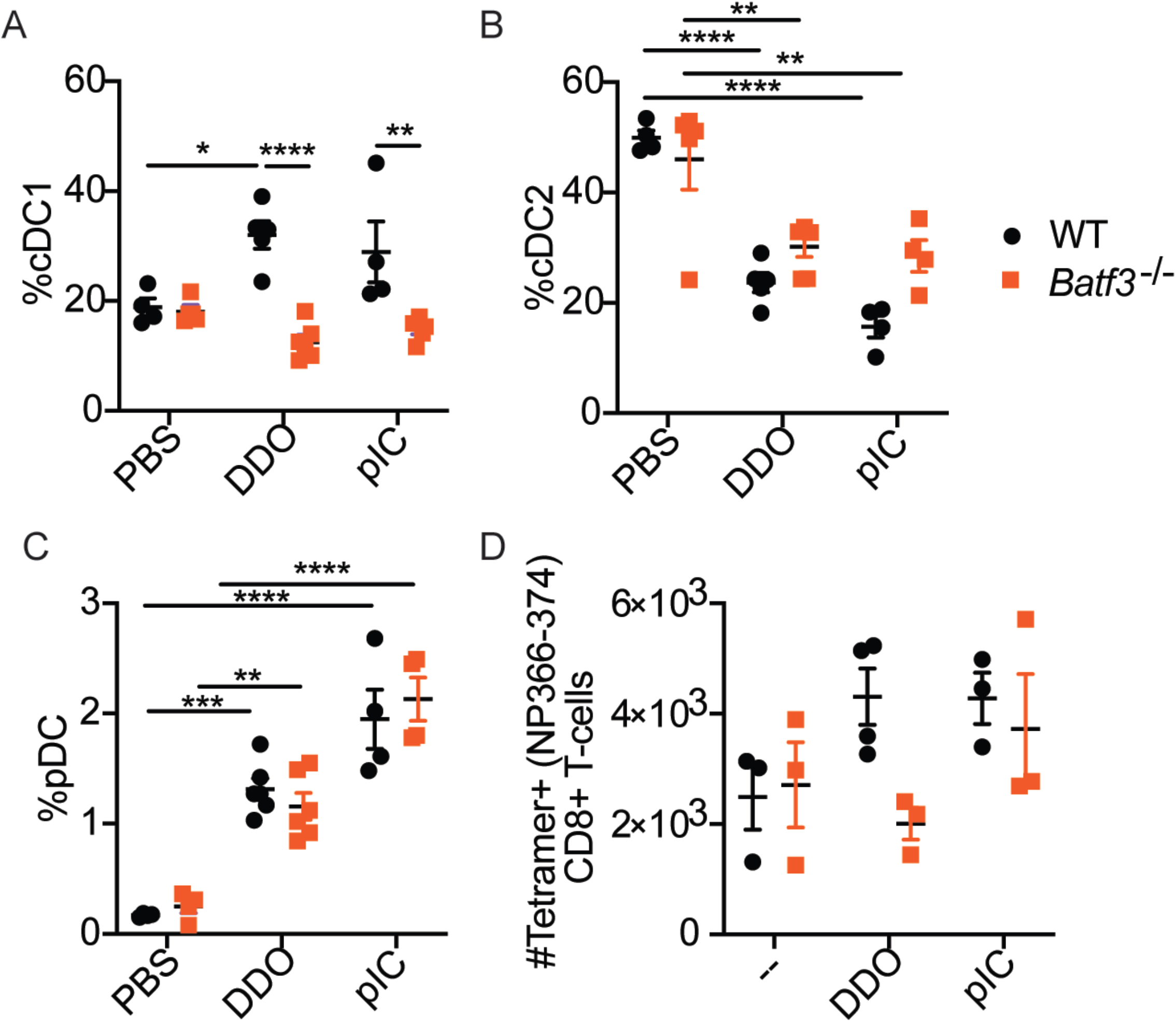
DDO requires cDC1s to induce a CD8+ T-cell response. **A-C** WT (black) and Batf3^-/-^ (orange) mice were injected s.c. with PBS, 5µg DDO, or 5µg poly I:C (pIC). Draining lymph node were harvested 12h post-injection and analyzed by flow cytometry (n=4-6/group). Mean±SEM of each group is shown. **A** Percent of conventional DCs that are cDC1s characterized as live, CD3^-^NK1.1^-^B220^-^CD19^-^, MHCII^hi^, CD64^-^,Ly6c^-^, CD11c^hi^, XCR1^+^SIRPa^-^ **B** Percent of conventional DCs that are cDC2s characterized as live, CD3^-^NK1.1^-^B220^-^CD19^-^,MHCII^hi^, CD64^-^,Ly6c^-^, CD11c^hi^, XCR1^-^SIRPa^+^. **C** Percent of cells in the lymph nodes that are pDCs were characterized as live, PDCA1^+^B220^+^. **D** WT and Batf3^-/-^ were immunized twice, 14d apart, with 10µg disIAV alone (--) or adjuvanted with 5µg DDO or 5µg pIC. Spleens were harvested 7d post-boost, processed into a single cell suspension, and analyzed by flow cytometry (n=3-4/group). Mean±SEM of each group is shown. **D** Number of tetramer^+^ CD8+ T-cells were defined as Live, CD3^+^, CD8^+^CD4^-^, tetramer^+^. **=p<0.01, ***=p<0.001, ****=p<0.0001. Data represent one representative experiment out of 2 independent repeats.

### 3.4 DDO induces a local type I IFN response

To investigate the early response to DDO injection, which leads to the accumulation of cDC1s in the draining lymph node, mice were injected s.c. with PBS, DDO, or poly I:C and the injected footpad and draining lymph node were harvested at 4, 12, and 24h post-injection for cytokine expression analysis by RT-qPCR. At 4h post-injection, *Ifnb1* transcripts peaked in the footpad and quickly returned to near baseline levels for both DDO and poly I:C (Figure 4A). Transcription of IFN-stimulated genes (ISGs), such as *Mx1* and the chemokine *Ccl5* peaked at 12h post-injection for both DDO and poly I:C (Figure 4B-C). *Mx1* returned to baseline expression levels by 24h but *Ccl5* remained elevated. Both DDO and poly I:C induced transient *Il6* expression at 4h post-injection (Figure 4D). Interestingly, DDO induced greater *Il1b* expression than poly I:C in the injection site, indicating differences in the inflammatory response between these RNA molecules (Figure 4E).

**Figure 4:**
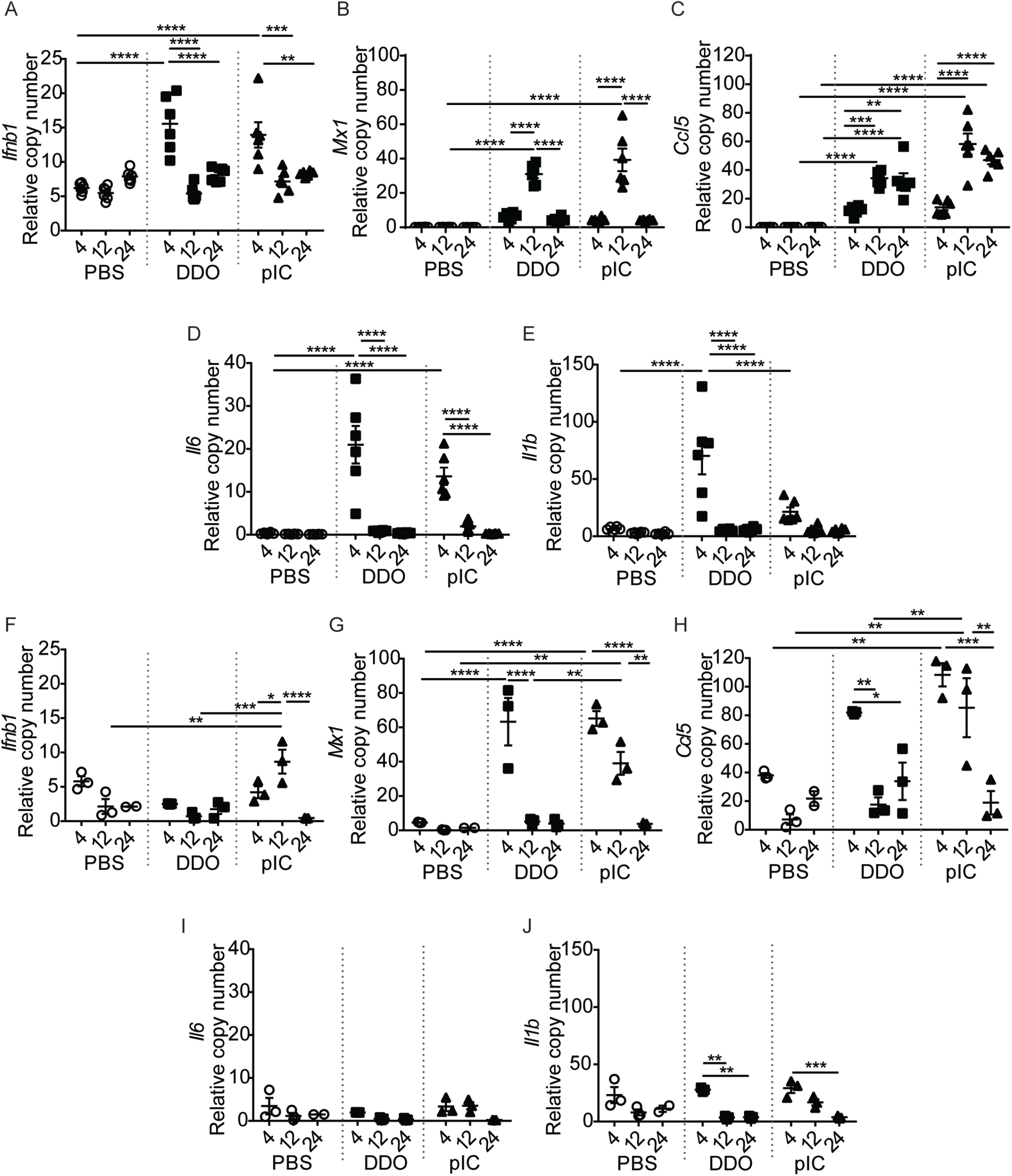
DDO activates a rapid type I IFN response with a different cytokine profile than poly I:C. WT mice were injected s.c. with PBS, 5µg DDO, or 5µg poly I:C (pIC) in the rear footpad. Injected footpads and draining lymph node were harvested 4, 12, and 24h post-injection. **A-E** Expression of transcripts in the footpad are relative to the housekeeping gene *β-actin*. (n=6/group). Mean±SEM of each group is shown. **A** Relative *Ifnb1*. **B** Relative *Mx1*. **C** Relative *Ccl5*. **D** Relative *Il6*. **E** Relative *Il1b*. **F-J** Two lymph nodes/group were pooled into one sample for better RNA quality. Expression of transcripts are relative to the housekeeping gene *β-actin*. (n=3/group). Data correspond to mean±SEM of each group. **F** Relative *Ifnb1*. **G** Relative *Mx1*. **H** Relative *Ccl5*. **I** Relative *Il6*. **J** Relative *Il1b*. *=p<0.05, **=p<0.01, ***=p<0.001, ****=p<0.0001. Data represent one representative experiment out of 2 independent repeats.

In the draining lymph node, greater differences in response to treatments were revealed. Only injection with poly I:C resulted in detectable *Ifnb1* transcripts in the draining lymph node at any time point (Figure 4F). Both DDO and poly I:C injection induced ISG transcription at 4h in the draining lymph node, however, poly I:C induced sustained ISG expression that lasted at least 12h post-injection (Figure 4G-H). These data support the induction of systemic type I IFN responses by poly I:C, a response not observed in DDO-injected mice. Inflammatory cytokines *Il6* and *Il1b* were not significantly induced over PBS injected mice in either condition, indicating a primarily local inflammatory response at these time points (Figure 4 I-J).

In both the footpad and the lymph node, other type I IFN gene transcripts were examined including *Ifna1, Ifna2, Ifna4, Ifna5, Ifna6, Ifna12*, and *Ifna13*. Both DDO and Poly I:C elicited small increases in *Ifna4* at 4 hr in the footpad. However, unlike Poly I:C, DDO does not induce any upregulation of any of the *Ifna genes examined* in the draining lymph node (Figure S1). As *Ifnb1* transcript copy numbers were the only interferon copy numbers that were elevated at the injection site, we determined that IFNβ was the primary IFN produced at the site of injection.

Together, these data show that DDO induces a local type I IFN response, in contrast to poly I:C, a known inducer of systemic type I IFN responses[22]. In addition to the magnitude of IFN responses, DDO induced a stronger expression of inflammatory cytokines than poly I:C. These data underscore the differences between the local responses to two seemingly similar RNAs, one a natural virus-derived pathogen molecular pattern (PAMP) and one a synthetic dsRNA, confirming previous observations with a prototype DDO at a single time point[13].

### 3.5 Type I IFN is required to initiate immune responses to DDO-adjuvanted vaccines

Type I IFN is a cytokine family with broad functions that include preparing cells for a viral challenge and helping to shape the adaptive immune response. Type I IFN can aid in the activation of DCs[23, 24] and lead to the activation and type 1 immunity polarization of T-cells[25, 26]. As we have observed a role for type I IFN in inducing T-cell responses upon i.m. immunization with DDO-adjuvanted vaccines, we next tested if type I IFN was necessary for cDC1 accumulation in the lymph node upon s.c. immunization. WT mice and mice deficient in type I IFN signaling (*Ifnar*^-/-^) were injected in the footpad with PBS or DDO and the draining lymph nodes were harvested at 12h post-injection to analyze the DC composition. In contrast to WT mice, upon DDO injection, there was no increase in the percentage of cDC1s in the draining lymph node in *Ifnar*^-/-^ mice (Figure 5A-B) indicating a dependence on type I IFN signaling for the accumulation of cDC1s in the draining lymph node. Additionally, there was no increase in the percentage of pDCs in the draining lymph nodes of DDO *Ifnar*^*-/-*^ mice (Figure 5C).

**Figure 5:**
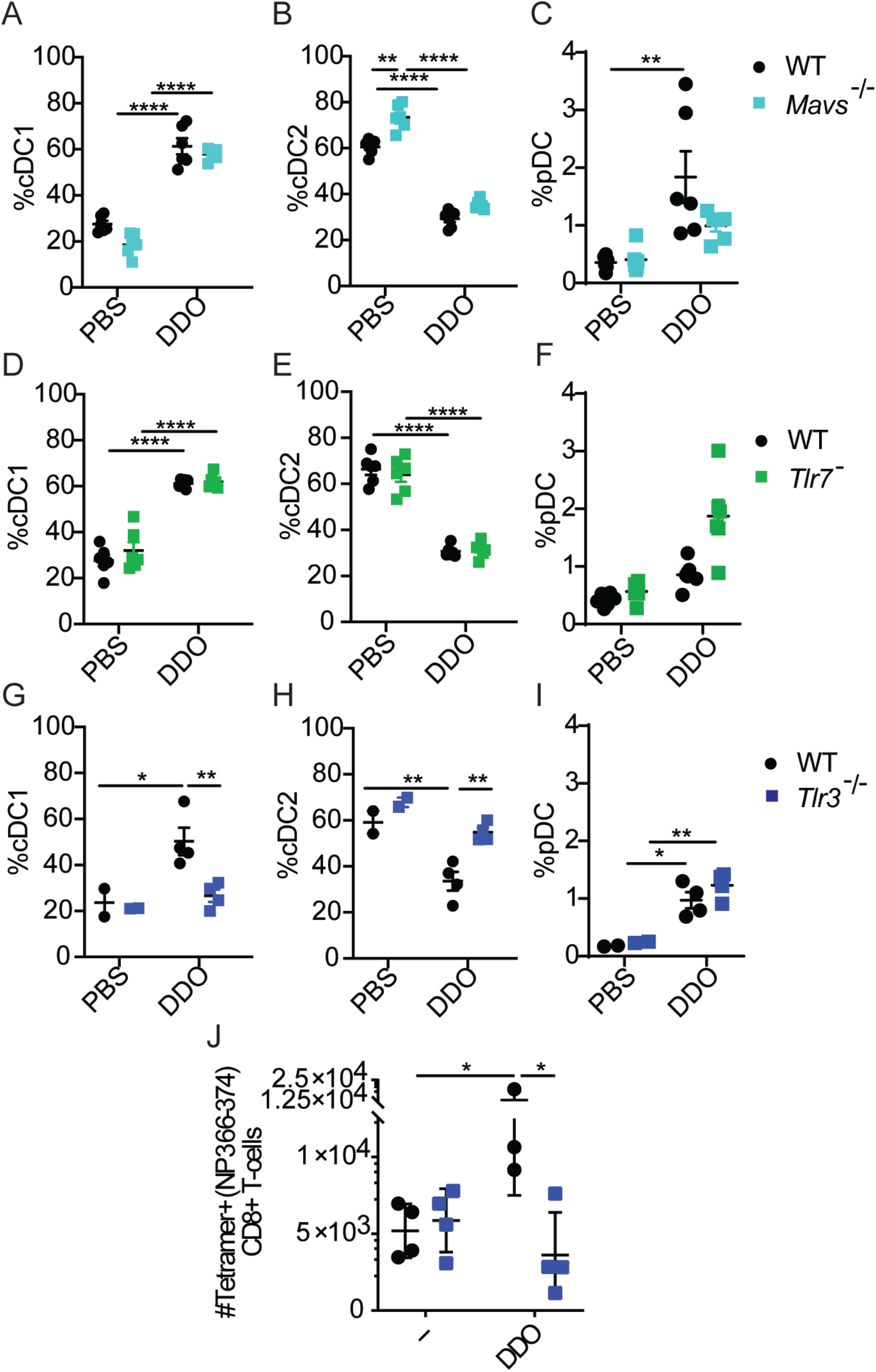
Type I IFN is required for cDC1 accumulation and the subsequent T-cell response to DDO-adjuvanted vaccines. WT (black) and *Ifnar*^-/-^ (red) mice were injected s.c. with PBS or 5µg DDO in the rear footpad. Draining lymph nodes were harvested 12h post-injection and processed into a single cell suspension for analysis by flow cytometry. (n=4-6/group) Mean±SEM of each group is shown. **(A-C). A** Percent of conventional DCs that are cDC1s characterized live, CD3^-^NK1.1^-^B220^-^CD19^-^, MHCII^hi^, CD64^-^, Ly6c^-^, CD11c^hi^, XCR1^+^SIRPa^-^. **B** Percent of conventional DCs that are cDC2s characterized as live, CD3^-^NK1.1^-^ B220^-^CD19^-^, MHCII^hi^, CD64^-^, Ly6c^-^, CD11c^hi^, XCR1^-^SIRPa^+^. **C** Percent of cells in the lymph nodes that are pDCs that are characterized as live, PDCA1^+^B220**^+^. D** WT and *Ifnar*^-/-^ mice were immunized twice, 14d apart, with 10µg disIAV alone (--) or adjuvanted with 5µg DDO. Spleens were harvested 7d post-boost, processed into a single cell suspension, and analyzed by flow cytometry (n=3-4/group). Mean±SEM of each group is shown. **D** Number of tetramer^+^ CD8+ T-cells were defined as Live, CD3^+^, CD8^+^CD4^-^, tetramer^+^. **=p<0.01, ****=p<0.0001. Data represent one representative experiment out of 2 independent repeats.

To determine if the dependence on type I IFN signaling for DC accumulation resulted in a reduction in CD8+ T-cell responses in this system, mice were vaccinated against IAV as described above. *Ifnar*^*-/-*^ mice vaccinated with DDO-adjuvanted vaccines were unable to generate IAV-specific CD8+ T-cells (Figure 5D), indicating a complete reliance on type I IFN for response to DDO-adjuvanted vaccines.

### 3.6 TLR3 is required for the accumulation of DCs in the draining lymph node and the subsequent CD8+ T-cell response in response to DDO

Adjuvants, including DDO and poly I:C, act as PAMPs[27]. PAMPs are sensed through various pattern recognition receptors (PRRs) including endosomal RNA sensors TLR7[28] and TLR3[27, 29] and cytosolic RNA sensors RIG-I and MDA5 which signal through the common adaptor MAVS for the production of type I IFN and other cytokines[30-33]. As type I IFN is required for both the accumulation of cDC1s in the draining lymph node and the adaptive immune response induced by DDO, we chose to examine how DDO is sensed upon s.c injection.

To investigate which of these sensors are responsible for the recognition of DDO and its immunostimulatory activity, WT mice and *Tlr7*^-^, *Tlr3*^-/-^, and *Mavs*^-/-^ mice were injected with PBS or DDO and DC composition in the lymph node was evaluated at 12h post-injection. WT and *Mavs*^-/-^ mice had similar changes in the DC composition after DDO injection (Figure 6A-C). Similar to the response in *Mavs*^-/-^ mice, the DC composition in the lymph nodes of DDO-injected *Tlr7*^-^ mice mirrored the response of WT mice (Figure 6D-F). However, injection of *Tlr3*^-/-^ mice, resulted in a loss of the cDC response in DDO-injected mice compared to WT mice (Figure 6G-H). The pDC response remained intact in *Tlr3*^-/-^ upon DDO injection (Figure 6I), likely due to the low expression level of TLR3 by pDCs. These data illustrate the critical role TLR3 plays in sensing DDO to lead to the accumulation of cDC1s in the draining lymph node.

**Figure 6:**
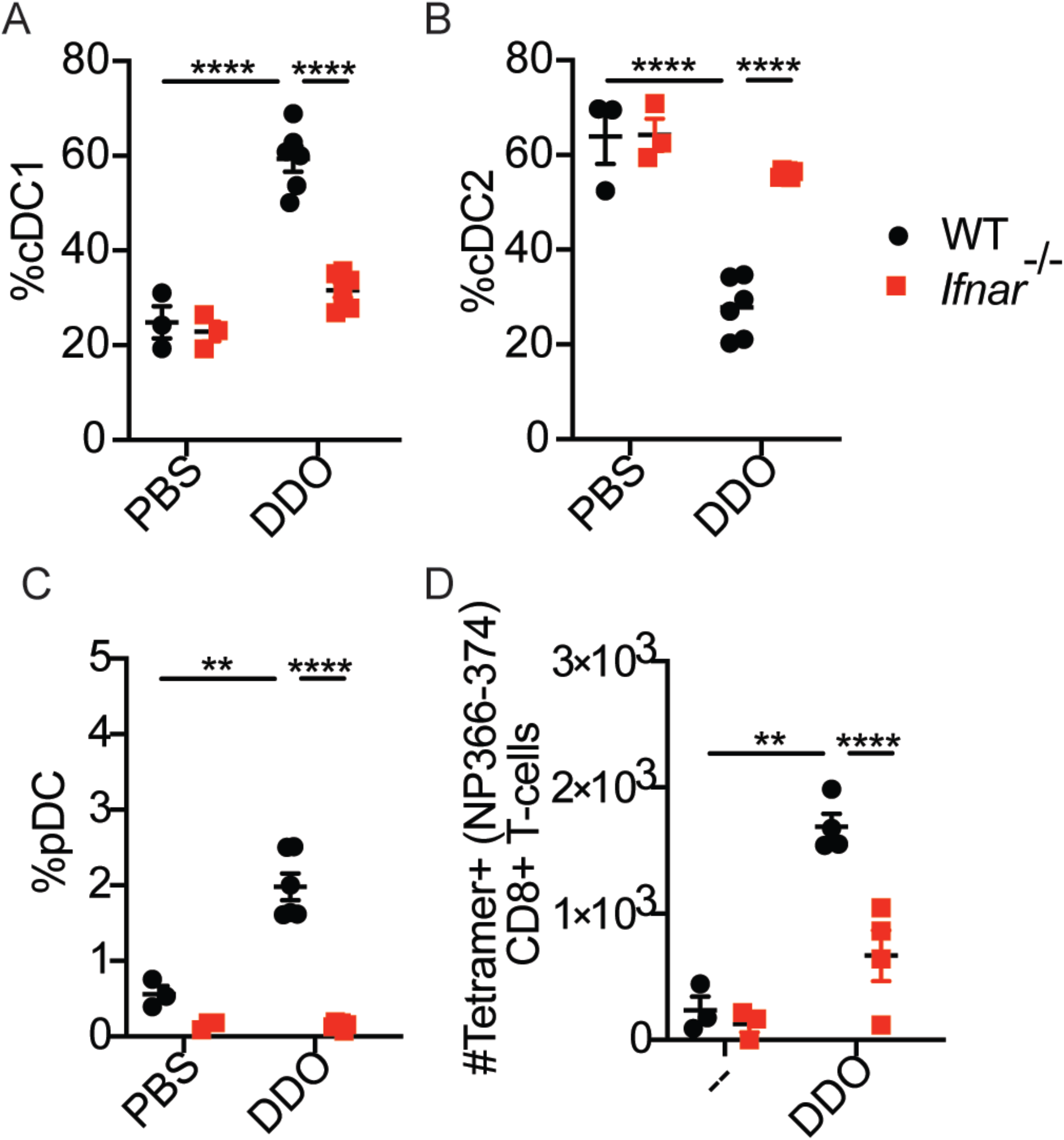
TLR3 is required for cDC1 migration in response to DDO injection. **A-C** WT (black) and *Mavs*^-/-^ (cyan) mice were injected s.c. with PBS or 5µg DDO in the rear footpad. Draining lymph nodes were harvested at 12h post-injection, processed into a single cell suspension, and analyzed by flow cytometry. (n=4-6/group) Mean±SEM of each group is shown. **A** Percent of conventional DCs that are cDC1s characterized live, CD3^-^NK1.1^-^B220^-^CD19^-^, MHCII^hi^, CD64^-^, Ly6c^-^, CD11c^hi^, XCR1^+^SIRPa^-^. **B** Percent of conventional DCs that are cDC2s characterized as live, CD3^-^NK1.1^-^B220^-^CD19^-^, MHCII^hi^, CD64^-^, Ly6c^-^, CD11c^hi^, XCR1^-^SIRPa^+^. **C** Percent of cells in the lymph nodes that are pDCs that are characterized as live, PDCA1^+^B220**^+^. D-F** WT (black) and *Tlr7*^-^ (green) mice were injected s.c. with PBS or 5µg DDO in the rear footpad. Draining lymph nodes were harvested at 12h post-injection, processed into a single cell suspension, and analyzed by flow cytometry. (n=6/group Mean±SEM of each group is shown. **D** Percent of conventional DCs that are cDC1s characterized as in A. **E** Percent of conventional DCs that are cDC2s characterized as in B. **F** percent of cells in the lymph nodes that are pDCs characterized as in C. **G-I** WT (black) and *Tlr3*^-/-^ (blue) mice were injected s.c. with PBS or 5µg DDO in the rear footpad. Draining lymph nodes were harvested at 12h post-injection, processed into a single cell suspension, and analyzed by flow cytometry. (n=2-4/group) Mean±SEM of each group is shown. **G** Percent of conventional DCs that are cDC1s characterized as in A. **H** Percent of conventional DCs that are cDC2s characterized as in B. **I** percent of cells in the lymph node that are pDCs characterized as in C. **J** WT and *Tlr3*^-/-^ mice were immunized twice, 14d apart, with 10µg disIAV alone (--) or adjuvanted with 5µg DDO. Spleens were harvested 7d post-boost, processed into a single cell suspension, and analyzed by flow cytometry (n=3-4/group). Mean±SEM of each group is shown. Number of tetramer^+^ CD8+ T-cells were defined as Live, CD3^+^, CD8^+^CD4^-^, tetramer^+^. *=p<0.05, **=p<0.01, ****=p<0.0001. Data represent one representative experiment out of 2 independent repeats.

As all conditions that abrogated cDC1 responses also eliminated T-cell responses to DDO-adjuvanted vaccines, we next examined the role of TLR3 during T-cell responses upon DDO-adjuvanted vaccinations. WT and *Tlr3*^-/-^ were vaccinated as above.

In concordance with *Ifnar*^-/-^ and *Batf3*^-/-^ studies, the loss of TLR3 and subsequent loss of cDC1 skewing in the lymph node resulted in a reduction of the antigen-specific CD8+ T-cell response. These data demonstrate a critical role for TLR3 in sensing DDO upon s.c. injections and inducing antigen-specific CD8+ T-cell responses.

## 4. DISCUSSION

Understanding the innate immune response needed to instruct a CD8+ T-cell response to vaccination is critical for targeted vaccine design. DDO is a 268nt RNA derived from the primary immunostimulatory molecule of SeV infections[12, 13]. Here we show that DDO induces a local type I IFN and inflammatory response that leads to the accumulation of cDC1s in the draining lymph node. cDC1 recruitment to the lymph node relies on TLR3 and type I IFN signaling and loss of cDC1 accumulation in the draining lymph nodes prevents the development of antigen-specific CD8+ T-cell responses.

TLR3 is a known poly I:C sensor[27] and we show here that it is also required for DDO-induced immune responses during vaccination in mice. However, the cytokine response to these PAMPs is not identical. DDO does not induce a systemic type I IFN response and triggers higher expression of *Il1b* than poly I:C. These intriguing results raise questions about how these differences arise and whether additional sensors or cell types are engaged by DDO at the injection site. Studies with licensed adjuvants, alum and MF59, have shown that additional innate cells such as natural killer cells and monocytes are critical for antigen transport to the lymph node[34]. Perhaps DDO and poly I:C interact with different cells to induce this differential cytokine response. Further examination of the exact cell types interacting with DDO is needed to fully understand its adjuvancy mechanism.

Another group using a full-length version of the SeV DVG (546nt) as a vaccine adjuvant showed that its ability to induce type I IFN and protect from lethal challenge upon i.m. injection is dependent on RIG-I[35]. We show that DDO, which is a truncated version of the full-length DVG[12], induced cDC1 accumulation in the draining lymph node and antigen specific CD8+T-cells in a TLR3-dependent manner, despite that DDO is sensed through RIG-I when transfected into cells, similar to the full length DVG[12]. It is possible that differences in the concentration of DDO used, DDO size, route of immunization, or time point analyzed explain these seemingly contradictory results.

Intriguingly, to trigger RIG-I, RNA would have to be internalized to the cytoplasm after injection, as is observed when the full-length DVG is injected[35]. Work from the Mossman lab has demonstrated the need for class-A scavenger receptors for the internalization and subsequent sensing of dsRNA through both TLR3 and RLRs[36, 37]. Additionally, proteins such as SIDT2 have been identified as RNA transporters from endosomes into the cytosol for RLR sensing[38]. Further studies are required to determine if class-A scavenger receptors or SIDT2 are used by DDO and if differences between the full-length DVG-adjuvanted vaccinations and DDO-adjuvanted vaccinations change how these related RNAs interact with proteins involved with RNA internalization and lead to sensing by TLRs or RLRs.

Our data show that DDO relies on cDC1s to induce CD8+ T-cell responses. Any loss of cDC1 accumulation in the draining lymph node, through loss of BATF3, type I IFN signaling, or TLR3, abrogated DDO’s adjuvancy. It is clear that cDC1s are a potent driver of CD8+ T-cell responses in this system and many viral infections, including West Nile virus[21], cytomegalovirus[39], influenza virus[40], cowpox virus[41], and others. However, the exact mechanism of cDC1 activation and PAMP sensing during many vaccinations is still unclear. Studies from the Kedl lab indicate the IFN-stimulated gene IL-27 production from cDC1s as an indicator of subsequent CD8+ T-cell responses after subunit vaccination[42, 43], however they did not examine the signals that lead to the activation of and IL-27 production by cDC1s. To gain a comprehensive picture of how CD8+ T-cell responses are induced, we must fully understand how the cells directly instructing CD8+ T-cells are activated.

We have shown DDO act as a type 1 immunity inducing adjuvant using an influenza vaccination model[11]. DDO was derived from a different virus than it was used to vaccinate against. It is likely that DDO would be beneficial in vaccines for other intracellular pathogens. Intracellular parasites remain a difficult target for vaccination and pathogens such as plasmodium, toxoplasma, and leishmania rely on type 1 immune responses for their clearance and control[44-46]. For effective immunizations to occur, the proper antigens must be targeted. Recent advances in epitope discovery, such as T-scan[47], are allowing for more targeted vaccine antigen design. By pairing appropriate antigens with DDO, tailored vaccinations against diverse pathogens could be tested.

Great strides have been made in understanding the difference between responses to infection and vaccination with a shift in the focus on the role of innate responses. We have shown an adjuvant derived from a natural PAMP is sensed by TLR3 and induces cDC1 accumulation in the draining lymph node and that this process is necessary for antigen specific CD8+ T-cell development. This study has uncovered the early innate immune response to DDO and has identified potential targets for type 1 immunity inducing adjuvants.

## AUTHORS CONTRIBUTIONS

Conceived experiments: D.G.F., and C.B.L.; performed experiments and collected data: D.G.F, D.J.H. Wrote the original draft: D.G.F. and C.B.L.; Supervised research activities: C.B.L.

## ACKNOWLEDGEMENTS

The authors wish to thank: Dr. Laurence Eisenlohr (Children’s Hospital of Philadelphia) for providing tetramer, Dr. Christopher Hunter and his lab (University of Pennsylvania) for providing mice and flow cytometry reagents, and Dr. David Christian (University of Pennsylvania) for advice on DC harvests and staining. This work was supported by the US National Institutes of Health National Institute of Allergy and Infectious Diseases (NIH R01 AI134862 to C.B.L).

## Figure Legends

**Figure S1:**
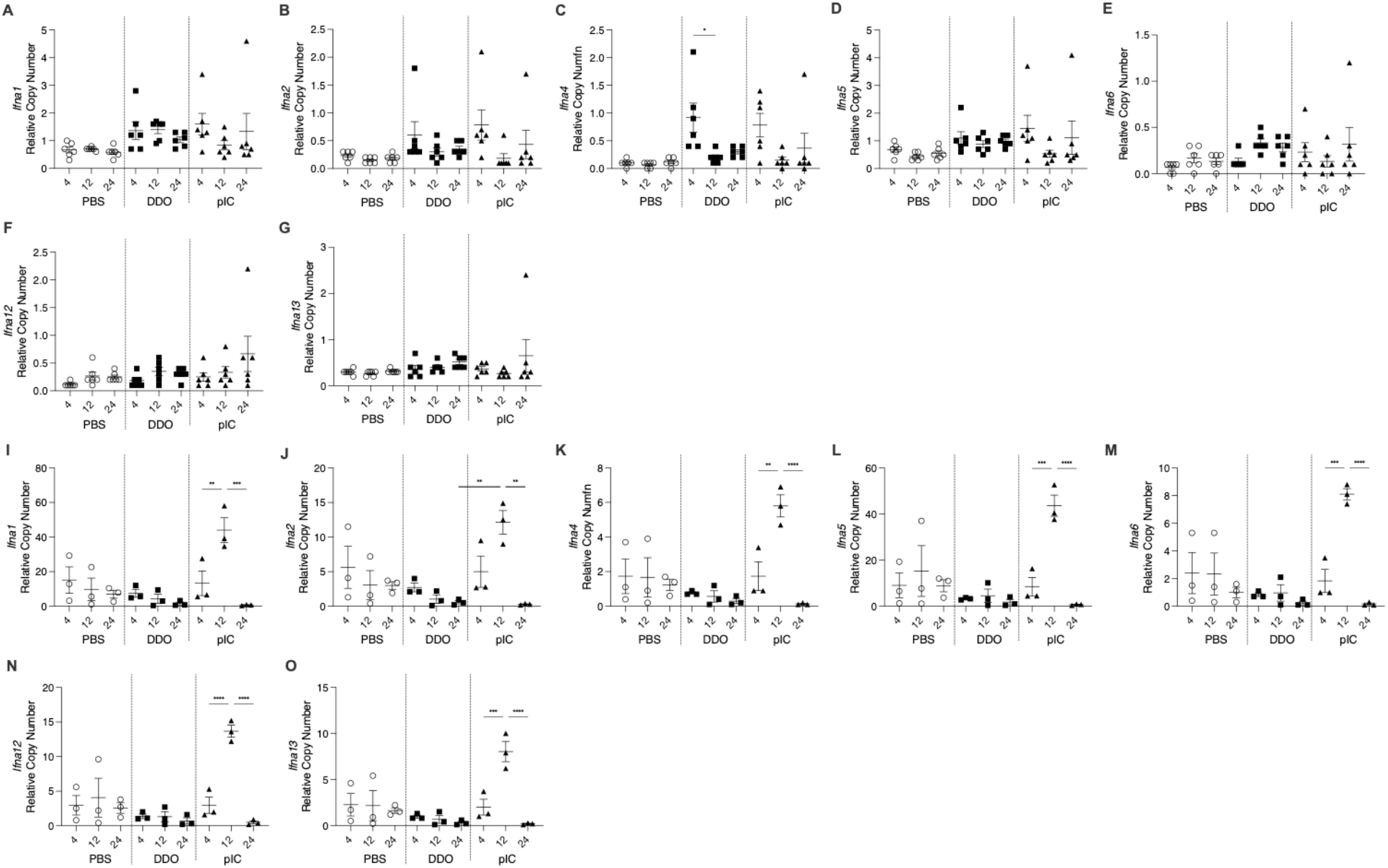
DDO does not induce a systemic IFNa response, in contrast to poly I:C. WT mice were injected s.c. with PBS, 5µg DDO, or 5µg poly I:C (pIC) in the rear footpad. Injected footpads and draining lymph node were harvested 4, 12, and 24h post-injection. **A-H** Expression of transcripts in the footpad are relative to the housekeeping gene *β-actin*. (n=6/group). Mean±SEM of each group is shown. Relative **A** *Ifna1*, **B** *Ifna2*, **C** *Ifna4*, **D** *Ifna5,* **E** *Ifna6*, **F** *Ifna12 anf* **G** *Ifna13.* **I-P** Two lymph nodes/group were pooled into one sample for better RNA quality. Expression of transcripts are relative to the housekeeping gene *β-actin*. (n=3/group). Data correspond to mean±SEM of each group. **I** Relative *Ifna1*, **J** *Ifna2*, **K** *Ifna4*, **L** *Ifna5,* **M** *Ifna6*, **N** *Ifna12,* **O** *Ifna13*, *=p<0.05, **=p<0.01, ***=p<0.001, ****=p<0.0001. Data represent one representative experiment out of 2 independent repeats.

